# Increased Osteoclast Activity Contributes to Bone Resorption and Osteopenia in a Rett Syndrome Mouse Model

**DOI:** 10.64898/2026.04.24.720567

**Authors:** Nadeem Samee, Lou Belz, Nicolas Narboux-Nême, Jean-Christophe Roux, Nicolas Panayotis, Giovanni Levi

## Abstract

Rett syndrome is a severe neurodevelopmental disorder caused predominantly by loss-of-function mutations in the X-linked gene *MECP2*. Besides a vast array of neurological and physiological impairments, patients also frequently develop severe osteopenia with increased fracture risk, however, the mechanisms underlying these skeletal defects are not completely understood. Previous work in *Mecp2*-null mouse models has suggested that osteopenia is mainly due to impaired osteoblast function and reduced bone formation. Here, we examined bone mass, microarchitecture, and remodeling parameters in a *Mecp2*-null mouse model during postnatal development, with a particular focus on osteoclast involvement. Micro-computed tomography and histomorphometric analyses showed reduced bone mineral density and trabecular bone volume, associated with increased trabecular separation and cortical thinning. These structural alterations were accompanied by increased osteoclast number per bone surface, elevated urinary deoxypyridinoline, and higher expression of osteoclast-associated genes, including *Cathepsin K*. Furthermore, gene expression analysis revealed an age-dependent shift in bone remodeling. At postnatal day 35, mutant mice showed reduced expression of *Dlx5* and *Dlx6*, consistent with low bone turnover. By postnatal day 55, *Rankl* and *Cathepsin K* were markedly upregulated, suggesting an increase in osteoclast resorptive activity, while key osteoblast markers and the RANKL/OPG ratio did not change significantly. A potential cell-autonomous contribution of *Mecp2* to osteoclast maturation is also suggested by the analysis of public transcriptomic datasets on human osteoclast differentiation. Together, our findings identify increased osteoclast activity as a significant contributor to Rett-associated osteopenia and suggest that skeletal pathology in *Mecp2* deficiency progresses from an early low-turnover state to a later phase of increased osteoclast resorption.

**HIGHLIGHTS:**

1. What are the main findings.

- *Mecp2*-null mice display reduced bone mass and altered bone microarchitecture during postnatal development, associated not only with reduced osteoblast activity, but also with increased osteoclast number, elevated urinary deoxypyridinoline, and increased expression of osteoclast-associated genes.
- Bone remodelling shows an age-dependent shift in *Mecp2* deficiency, from an early low-turnover state at postnatal day 35 to increased osteoclast resorptive activity at postnatal day 55.
2. What are the implications of the main findings?

- Rett-associated osteopenia is not explained solely by impaired osteoblast function, but also involves a significant osteoclast contribution to skeletal deterioration.
- These findings refine the pathophysiological model of bone involvement in Rett syndrome and support the idea that skeletal alterations evolve dynamically during disease progression.

## INTRODUCTION

Rett syndrome (RTT) is a severe neurodevelopmental disorder that predominantly affects girls, with an incidence of approximately 1 in 10,000 live births [1,2]. Following an initial period of apparently normal development, affected individuals undergo a progressive regression of motor and cognitive functions, accompanied by stereotypic hand movements, breathing abnormalities, and gait disturbances. In more than 96% of cases, RTT is caused by loss-of-function mutations in the X-linked gene *MeCP2*, which encodes the methyl-CpG–binding protein MeCP2, a key regulator of transcriptional programs that binds preferentially to methylated CpGs [3–5]; some other rare genetic variants (e.g., CDKL5, FOXG1) may also present with Rett-like features [6]. Heterozygous *MECP2* mutations are, in general, lethal in males and lead to a poly-handicap in heterozygous females [7].

Although *MeCP2* expression is particularly abundant in the central nervous system, RTT is now recognized as a multisystem disorder affecting several peripheral tissues, including the skeletal system [8,9]. RTT patients frequently present reduced bone mineral density, osteopenia, scoliosis, and an increased incidence of fractures, often emerging early in life and worsening with age and reducing their quality of life [10,11]. Clinical studies have reported significantly reduced bone mineral content and density in RTT patients compared with age-matched controls, indicating that skeletal involvement is a consistent and clinically relevant component of the disease [12–14]. For example, the bone mineral density, bone mineral content and spinal mineral density were significantly reduced in a group of 20 RTT subjects when compared to age and weight matched controls [15]. Bone histomorphometry of the iliac crest of RTT patients showed that low bone volume was accompanied by low bone formation rates [16,17]. However, the cellular mechanisms underlying RTT-associated bone fragility remain incompletely understood.

Bone homeostasis is maintained through a tightly regulated balance between bone-forming osteoblasts and bone-resorbing osteoclasts. Disruption of this balance, either through impaired bone formation or excessive bone resorption, can lead to osteopenia and osteoporosis [18]. Several studies in RTT patients and mouse models have suggested that reduced bone formation contributes to skeletal defects. In particular, a comprehensive analysis of the *Mecp2*-null mouse model demonstrated impaired osteoblast function, reduced mineral apposition rates, and defective bone material properties, leading to the conclusion that decreased bone formation predominates in RTT-associated bone loss [11,19,20]. However, these studies primarily relied on formation indices and static assessments of osteoclast number, leaving open the question of whether osteoclast function and bone resorption may also be altered.

Importantly, osteoclast-mediated bone resorption can increase without major changes in osteoclast number or in classical coupling signals such as the RANKL/OPG ratio, particularly when alterations affect osteoclast activity, lifespan, or resorptive capacity rather than differentiation alone [21,22]. Thus, an exclusive focus on bone formation parameters or osteoclast counts may overlook functionally relevant changes in bone resorption that contribute to remodeling imbalance. In addition, bone remodeling evolves during postnatal development and disease progression, raising the possibility that osteoclast involvement in *Mecp2* deficiency may be age-dependent and therefore underappreciated in previous studies [13,18,22].

In this study, we re-examined the skeletal phenotype of *Mecp2*-null mice with a specific focus on bone remodeling and osteoclast function. By combining bone densitometry, microarchitectural analyses, histomorphometry, systemic markers of bone resorption, and gene expression profiling, we show that *Mecp2* deficiency is associated with increased osteoclast activity and enhanced bone resorption, revealing a remodeling imbalance that evolves during distinct postnatal stages. Our findings indicate that osteopenia in the *Mecp2*-deficient mouse model for RTT reflects a remodeling imbalance involving increased osteoclast-mediated resorption, highlighting a more complex pathophysiological mechanism than reduced bone formation alone.

## MATERIALS AND METHODS

### Animals

Experiments were performed on the B6.129P2(c)-Mecp2^tm1-1Bird^/J mouse model for RTT [23] carrying targeted disruption of *Mecp2*.

The mutation was maintained on the C57BL/6J genetic background. 35-days-old (P35) and 55-days-old (P55) hemizygous males (named *Mecp2*^-/yBIRD^ throughout the manuscript) and their wild-type (WT) littermates were used.

The experimental procedures were carried out in line with the European Union directive 2010/63/EU for the care and use of laboratory animals. Genotyping was performed by routine PCR technique according to the Jackson Laboratory protocols as previously described [23].

### Measurement of bone mineral density by Dual-energy X-ray absorptiometry (DXA)

Dual-energy X-ray absorptiometry (DEXA) analysis was carried out under general anesthesia. Total body, whole femur, and caudal vertebral bone mineral content (BMC, mg), bone area (area, cm2), and bone mineral density (BMD, mg/cm^2^) were measured using a PIXImus x-ray densitometer (Lunar, France; Software version 1.44) in ultrahigh resolution mode (resolution 0.18×0.18 mm). The precision and reproducibility of the instruments had previously been evaluated by calculating the coefficient of variation of repeated DXA measurements. The coefficient of variation was <2% for all the parameters evaluated. A calibration phantom was scanned daily to monitor the stability of the measurements.

### Histomorphometry and microcomputed tomography analysis

The left femur from each animal was excised at death, and the surrounding soft tissue was cleaned off. After storage in 70% ethanol at 4°C, the femurs were trimmed, and the distal halves of bones were post-fixed in 70% ethanol, dehydrated in xylene at 4°C, and embedded without demineralization in methyl methacrylate. Histomorphometric parameters were measured in accordance to the ASBMR nomenclature [24,25] on 5µm sections using a Nikon microscope interfaced with the software package Microvision Instruments (Evry, France). Sections were stained with aniline blue. For TRAP detection, sections were stained with a 50mM sodium tartrate and naphtol ASTR phosphate (Sigma, St Louis, France). The measurements of the trabecular bone were performed in a region of the secondary spongiosa. The trabecular bone volume (BV/TV), the trabecular bone thickness (Tb.Th), and the trabecular separation (Tb.Sp) and trabecular number (Tb.N) were measured. For cortical bone, we measured the average bone and marrow diameters at the femoral metaphysis and calculated the cortical thickness (Cort.Th).

The other femurs and tibias from 35 days- and 55 days-old *Mecp2*^-/yBIRD^ mice and sex and age matched control littermates were used for three dimensional (3D) microcomputed tomographic (μCT) analyses of the cortical thickness at the femoral midshaft. MicroCT analysis was realised by using a Skyscan 1072 scanner.

### Bone resorption markers in urine

Pyridinoline cross-links are released during the process of collagen breakdown and are cleared by the kidneys. In particular, Deoxypyridinoline (DPD) is released during bone resorption into the blood stream and is eliminated unmodified with urine. To quantify osteoclastic bone resorption, animals were fasted overnight, and urine samples were collected to measure the level of DPD cross-linkage using a chemiluminescent assay on an Immulite 2000 automated analyzer (DPC Siemens Medical Solutions Diagnostics, La Garenne-Colombes, France). Values are reported relative to creatinine concentrations (DPD/Cr) as determined by a standardized colorimetric assay using alkaline picrate with a Advia 2400 automated analyzer (Siemens Medical Solutions Diagnostics, Puteaux, France).

### RNA extraction and Real-time quantitative PCR (qPCR)

For long bone RNA preparations, soft tissues surrounding the bones were stripped off. The epiphyses were cut off and the bone marrow flushed out with PBS. Total RNAs were isolated using Trizol (Invitrogen, Carlsbad, CA) and processed with an RNeasy cleanup kit (Qiagen, Valencia, CA) according to the manufacturer’s instructions then reverse-transcribed into cDNA using the Reverse-iT Max Blend kit (ABgene, Surrey, UK). Quantitative real-time PCR expression analysis was performed on Lightcycler (Roche Diagnostics) using Absolute® SYBR Green capillary mix (ABgene) at 56°C for 40 cycles. Primers were designed using the online mouse library probes of Roche Diagnostics. mRNA levels were normalized against either Aldolase A or 18S.

### Statistical analysis

*In vitro* experiments were independently performed three times. Data are presented as mean ± standard error of the mean (SEM). A p-value < 0.05 was considered statistically significant. For non-qPCR datasets, statistical differences between genotypes were assessed using two-way ANOVA, as appropriate. Gene expression levels were quantified by real-time quantitative PCR and expressed as relative expression values using the 2^−ΔΔCt method, normalized to the selected housekeeping gene and referenced to the control condition [26]. Statistical analyses were conducted separately for each developmental stage, P35 and P55, to allow stage-specific interpretation of genotype effects. For each gene, differences between control and mutant groups were evaluated using a two-tailed Welch’s t-test, which does not assume equal variances. Because 2^−ΔΔCt values are log-normally distributed, statistical analyses were performed after log2 transformation, corresponding to analysis on the ΔΔCt scale [27]. Given the small number of biological replicates, formal assessment of normality was considered underpowered; therefore, qPCR results were interpreted together with effect size, direction of change, and biological consistency [28]. Gene selection was restricted to a predefined set of biologically relevant targets involved in osteoblast–osteoclast coupling; therefore, p-values were not adjusted for multiple comparisons, to avoid an excessive increase in type II error in this small, hypothesis-driven qPCR dataset.

## RESULTS

### *Mecp2* Deficiency Leads to Progressive Loss of Bone Mass

To evaluate the impact of *Mecp2* deficiency on bone mass, bone mineral density (BMD) was measured by dual-energy X-ray absorptiometry (DXA) in femora and tibiae of *Mecp2*^-/yBIRD^ mice and wild-type littermates at postnatal day 35 (P35) and postnatal day 55 (P55). As previously reported [11,19,23], *Mecp2*-deficient mice exhibited reduced body size and weight compared with wild-type controls, but appeared healthy and did not display major locomotor impairments.

DXA analysis revealed a marked reduction in femoral BMD in *Mecp2*^-/yBIRD^ mice at P55 compared with wild-type littermates (−29%, *p* < 0.001; *n* = 4 per genotype; Figure 1A). At P35, differences in femoral BMD were smaller and did not reach statistical significance. A similar trend was observed in the tibia at both ages (Figure 1B). These findings indicate that *Mecp2* deficiency results in a progressive reduction in bone mass, with skeletal deficits becoming more pronounced with age.

**Figure 1.**
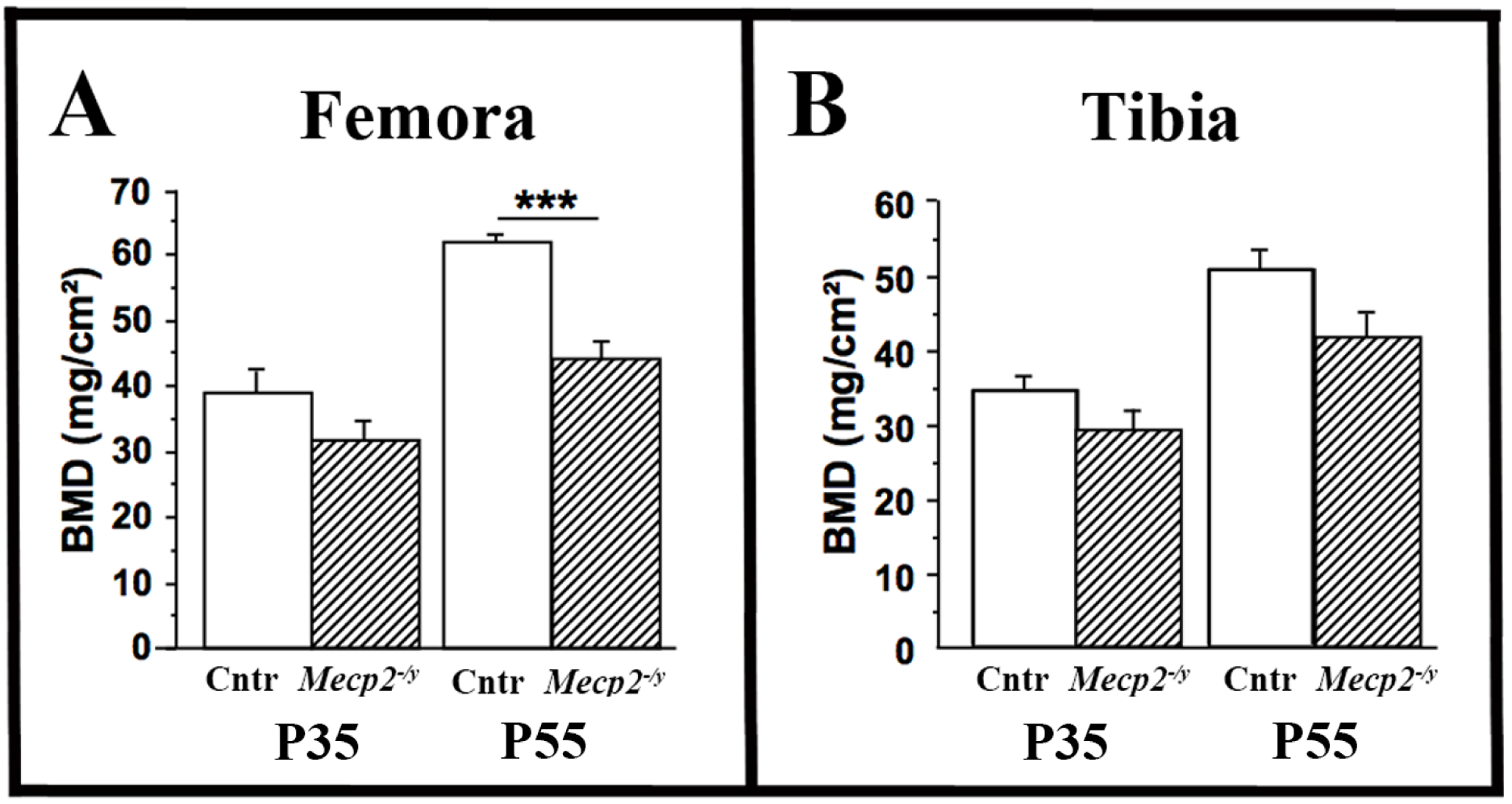
Effects of *Mecp2* deficiency on femoral BMD. BMD of 35- and 55-days-old (noted respectively P35 and P55) wild-type mice (open bars) and *Mecp2^−/yBIRD^*mice (hatched bars) were evaluated on the femurs by DXA. Results are presented as mean ± SEM; \*\*\**p* < 0.001; n=4 per genotype in all cases.

### *Mecp2* Deficiency Impairs Trabecular Bone Microarchitecture

To further characterize skeletal alterations, three-dimensional microcomputed tomography (µCT) was used to assess femoral bone microarchitecture. At P55, *Mecp2*^-/yBIRD^ mice displayed a significant reduction in total femoral bone volume (BV/TV) compared with wild-type controls (BV/TV: −35%, *p* < 0.05; *n* = 4 per genotype; Figure 2B), whereas no significant differences were detected at P35.

**Figure 2.**
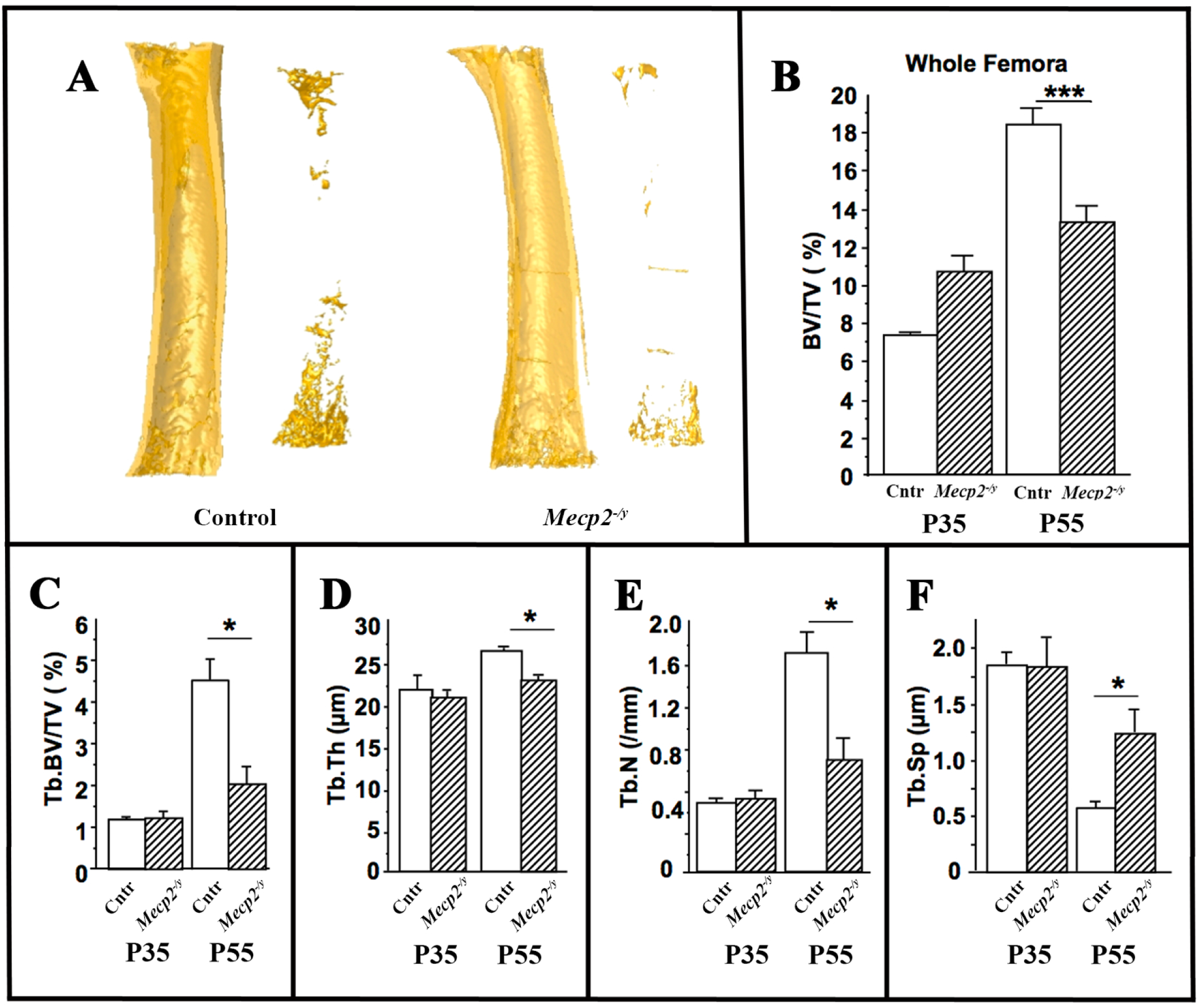
Impaired bone microarchitecture in Mecp2-null mice. Microarchitecture of femoral bone from wild-type mice (open bars) and *Mecp2 ^−/yBIRD^* mice (hatched bars) at P35 and P55 analyzed by µCT. A) Representative example of femurs from P55 WT and *Mecp2 ^−/yBIRD^* mice. Total bone volume (B) of the whole femur (BV/TV; %), trabecular bone volume (C) (Tb. BV/TV, %) and trabecular thickness (D) (Tb.Th), trabecular number (E) (Tb.N) and trabecular separation (F) (Tb.Sp) were measured. Mean ± SEM; (n=4 per group); \**p* < 0.05 and \*\*\**p* < 0.001.

Analysis of trabecular bone at the distal femoral metaphysis revealed pronounced microarchitectural deterioration in *Mecp2*-deficient mice at P55. Trabecular bone volume (Tb.BV/TV) was reduced by 55% (*p* < 0.05, Figure 2C), accompanied by significant decreases in trabecular thickness (Tb.Th: −12%, *p* < 0.05, Figure 2D) and trabecular number (Tb.N: −48%, *p* < 0.05, Figure 2E), and a marked increase in trabecular separation (Tb.Sp: +124%, *p* < 0.05, Figure 2F). No significant trabecular alterations were detected at P35. This pattern of trabecular deterioration is characteristic of pathological bone loss associated with altered bone remodeling dynamics with a possible decrease in osteoblast activity and increased osteoclasts activity.

### Cortical Bone Thickness Is Reduced in Symptomatic Mecp2-Deficient Mice

Cortical bone parameters were analyzed by µCT at the femoral midshaft. At P55, *Mecp2*^-/yBIRD^ mice exhibited a significant reduction in cortical thickness compared with wild-type littermates (−16%, *p* < 0.01; *n* = 4 per genotype; Figure 3A). This reduction resulted from concomitant decreases in both external diameter and marrow diameter (Figure 3B and C), indicating altered cortical modeling dynamics. No significant differences in cortical parameters were observed at P35. These results demonstrate that during postnatal growth cortical bone is progressively affected in *Mecp2*-deficient mice, consistent with an age-dependent skeletal phenotype.

**Figure 3.**
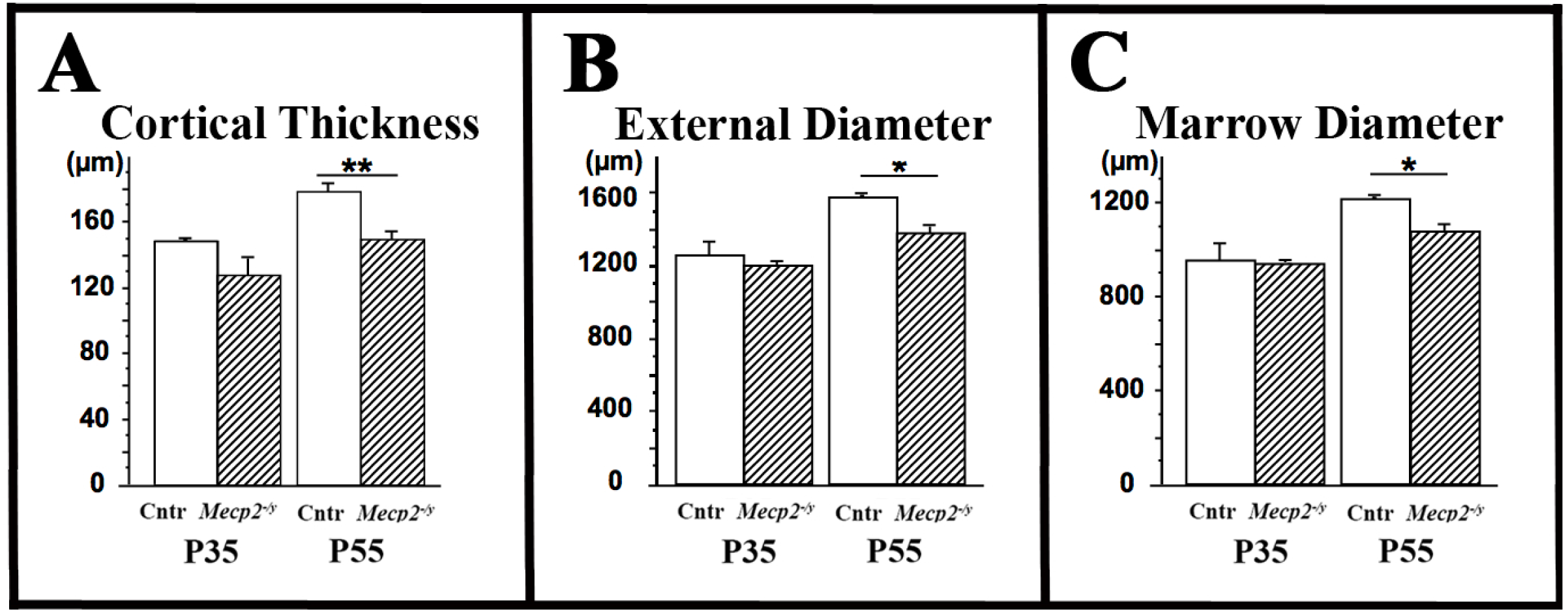
µCT analysis of cortical bone thickness of *Mecp2 ^−/yBIRD^* mice. µCT analysis of the femoral thickness at midshafts of 35- and 55-days-old (noted respectively P35 and P55) wild-type mice (open bars) and *Mecp2 ^−/yBIRD^* mice (hatched bars). Values of cortical thickness (A), external diameter (B) and marrow diameter (C) are presented as mean ± SEM; (n=4 per group). \**p* < 0.05 and \*\**p* < 0.01.

### Increased Osteoclast Number and Bone Resorption in Mecp2-Deficient Mice

To directly assess bone remodeling, histomorphometric analyses were performed on distal femora. At P35, trabecular structure and osteoclast parameters were comparable between *Mecp2*^-/yBIRD^ mice and wild-type controls. In contrast, at P55 *Mecp2*-deficient mice exhibited a significant reduction in trabecular bone volume (−42%, *p* < 0.05, Figure 4A), trabecular thickness (−17%, *p* < 0.05, Figure 4B), and trabecular number (−33%, *p* < 0.01, Figure 4C), accompanied by increased trabecular separation (+64%, *p* < 0.01, Figure 4D).

**Figure 4.**
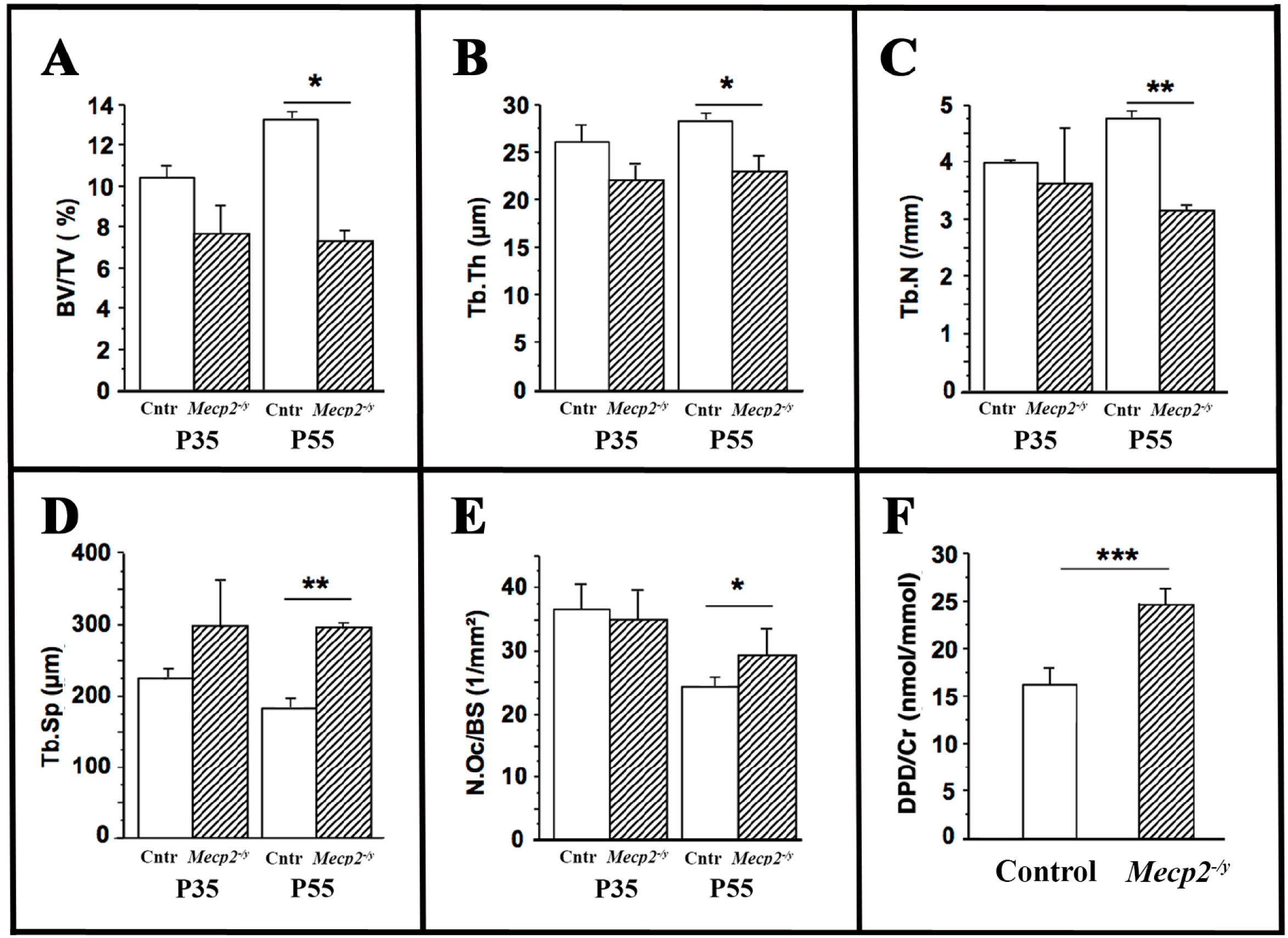
Histomorphometrical analysis: Increased number of osteoclasts in *Mecp2*^-/yBIRD^. Histomorphometry analysis was performed on distal femora of wild-type (open bars) and *Mecp2 ^−/yBIRD^* mice (hatched bars) at P35 and P55. The trabecular bone volume (A) (BV/TV, %) trabecular thickness (B) (Tb.Th), trabecular number (C) (Tb.N) and trabecular separation (D) (Tb.Sp) were measured. TRAP positive cells were counted on the trabecular surfaces of femurs. Results are presented as number of active osteoclasts per bone surface (N.Oc/BS) (E). Levels of urinary deoxypyridinoline/creatinine ratio (DPD/Cr) at P55 (F). Values of are presented as mean ± SEM; (n=4 per group); \**p* < 0.05; \*\**p* < 0.01, \*\*\**p* < 0.001.

Importantly, the number of osteoclasts per bone surface (N.Oc/BS) was significantly increased in P55 *Mecp2*^-/yBIRD^ mice (*p* < 0.05; Figure 4E), whereas osteoclast parameters were unchanged at P35. To determine whether increased osteoclast number was associated with enhanced resorptive activity, systemic bone resorption was assessed by measuring urinary deoxypyridinoline/creatinine (DPD/Cr) ratios [29]. *Mecp2*-deficient mice displayed a robust increase in DPD/Cr levels at P55 (+38%, *p* < 0.001; *n* = 4 per genotype; Figure 4F), indicating elevated osteoclast-mediated collagen degradation.

Together, these results demonstrate that osteoclast activity is increased in *Mecp2*-deficient mice at symptomatic stages, coinciding with the onset of pronounced bone loss.

### Expression of osteoblastic and osteoclastic markers in long bone of *Mecp2^-/yBIRD^*

To further characterize the molecular basis of the skeletal phenotype, the expression of osteoblast- and osteoclast-associated genes was analyzed in long bones by quantitative PCR at P35 and P55 (Figure 5). The expression levels of the osteoblastic markers *Runx2* and *osteocalcin* were not significantly altered at either age, suggesting that the skeletal phenotype was not associated with major changes in osteoblast differentiation. In contrast, *Mecp2*^-/yBIRD^ long bones showed a marked reduction in the level of *Dlx5* and *Dlx6* expression at P35, but not at P55. As *Dlx5* and *Dlx6* are transcription factors involved in osteoblast function and in the control of osteoblast/osteoclast coupling [30] their early reduction is consistent with an initial alteration of bone remodelling. At P55, the expression levels of both *Rankl* and *Opg* were significantly increased, whereas the *Rankl/Opg* ratio did not differ significantly between genotypes. The osteoclastic marker Tartrate-Resistant Acid Phosphatase (*Trap*) was unchanged both at P35 and at P55, while the expression of *Cathepsin K,* a protease involved in the control of bone resorption, was strongly increased at P55. Together, these findings suggest an age-dependent shift in bone remodelling in *Mecp2*^-/yBIRD^ mice, from an early molecular alteration compatible with low bone turnover to a later stage associated with increased osteoclast resorptive activity.

**Figure 5.**
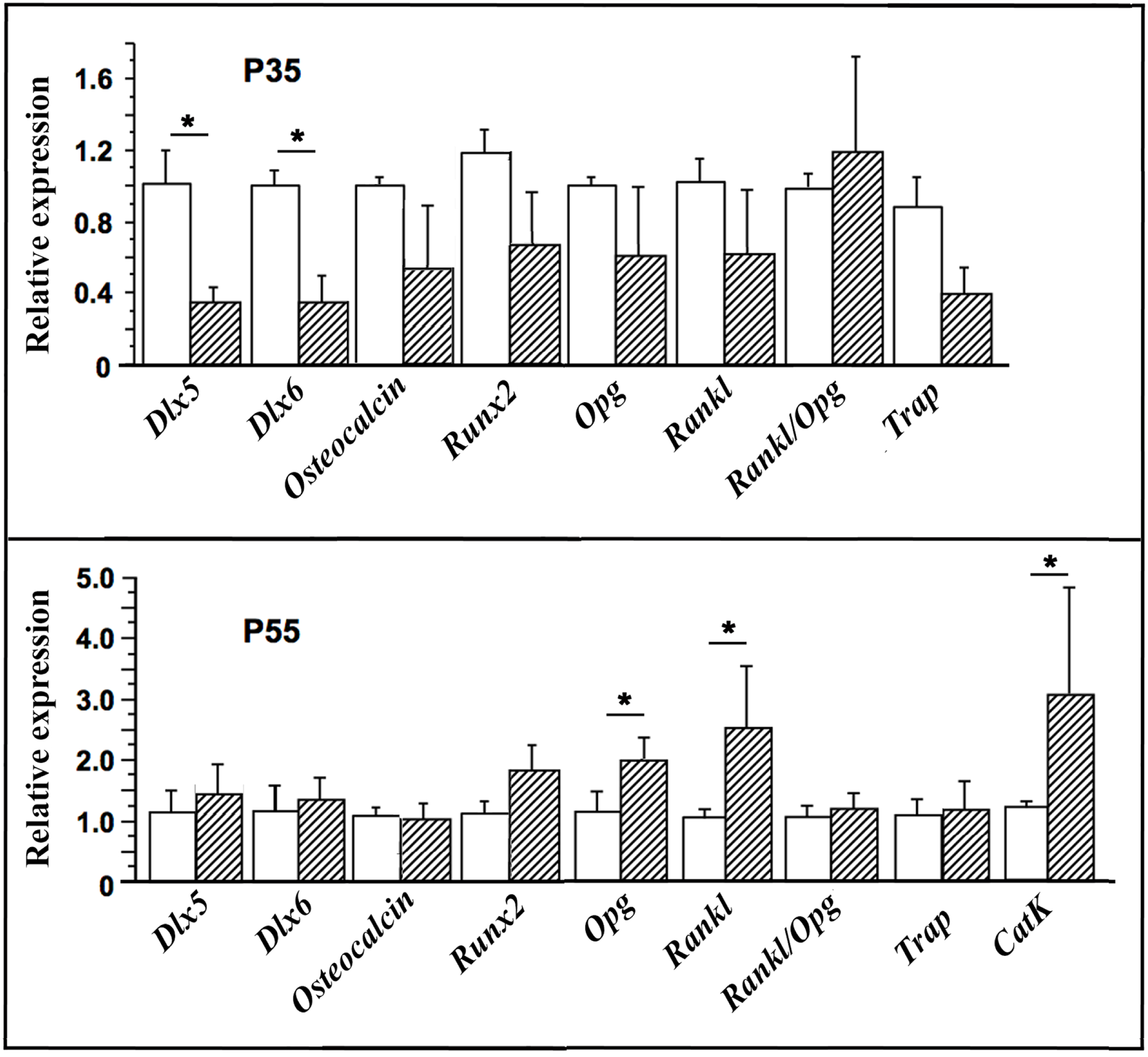
Relative expression levels of osteoblast- and osteoclast-related markers in wild type and *Mecp2 ^−/yBIRD^* mice femurs. The relative expression levels of *Dlx5*, *Dlx6*, *Osteocalcin, Runx2*, *Opg*, *Rankl*, *Trap* and *Cathepsin K* were measured by qPCR in wild type (open bars) and *Mecp2 ^−/yBIRD^* (hatched bars) littermates at P35 (upper graph) and P55 (lower graph). While the levels of *Dlx5* and *Dlx6* expression were reduced in *Mecp2 ^−/yBIRD^*mice at P35, *Osteocalcin, Runx2*, *Opg* and *Rankl* were unchanged and *Trap* expression was not significantly reduced. *Cathepsin K* expression was higher at P55. Although *Opg* and *Rankl* expression levels were increased at P55, the Rankl/Opg ratio did not change significantly at both ages studied. \**p* < 0.05.

## DISCUSSION

Although MeCP2 is best known for its essential role in neuronal function and synaptic maintenance, the frequent occurrence of osteopenia, scoliosis, and fractures in RTT indicates that MeCP2 deficiency also affects skeletal homeostasis. In the present study, we re-examined the bone phenotype of the *Mecp2*^-/yBIRD^ mice [23] and found evidence that bone loss in this model cannot be explained solely by impaired bone formation, as previously proposed, but also involves increased osteoclast-mediated resorption.

Our histological and microarchitectural analyses confirm that *Mecp2* deficiency is associated with reduced total and trabecular bone volume, decreased trabecular number, increased trabecular separation, and cortical thinning. These changes are consistent with the osteopenic phenotype described in RTT patients [11] and in previous analyses of the same mouse model [11]. Earlier studies concluded that bone loss in RTT predominantly reflects reduced bone formation, based largely on decreased mineral apposition rates and altered osteoblast function. Our data support a more complex view in which altered bone formation and increased bone resorption contribute in a temporally distinct manner to skeletal deterioration.

A major result of this study is that the molecular signature of bone remodelling changes with age. At P35, *Mecp2*^-/yBIRD^ mice showed reduced expression of *Dlx5* and *Dlx6*, consistent with an early low-turnover state. This pattern suggests that bone remodelling is globally reduced at this stage, rather than simply shifted toward excessive resorption. By contrast, at P55, expression of *Rankl* and *Cathepsin K* was significantly increased, whereas *Trap* expression and the *Rankl/Opg* ratio were not significantly altered. Together with the increased osteoclast number per bone surface and the marked elevation of urinary DPD/Cr ratio, these findings indicate that the later skeletal phenotype is associated with enhanced osteoclast resorptive activity.

This point is important because osteoclast-mediated bone resorption may increase without major changes in osteoclast number or in classical coupling indices such as the RANKL/OPG ratio, particularly when the main alteration affects osteoclast activity or resorptive capacity rather than differentiation alone. In this respect, the strong increase in *Cathepsin K* at P55 is especially informative, as this protease is a major effector of matrix degradation by mature osteoclasts. The combination of increased *Cathepsin K* and unchanged *Trap* suggests that in *Mecp2*^-/yBIRD^ mice, bone loss is linked more closely to enhanced osteoclast function than to a broad increase in osteoclast differentiation.

In parallel, we did not detect significant changes in the expression of the canonical osteoblast differentiation markers *Runx2* and *osteocalcin* at either stage. This suggests that the observed skeletal phenotype is not associated with major transcriptional alterations in terminal osteoblast differentiation markers in whole bone. However, the reduction of *Dlx5* and *Dlx6* at P35 indicates that earlier regulatory changes affecting osteoblast-related pathways may still occur. These findings therefore argue against a simple model in which RTT-associated osteopenia results exclusively from defective osteoblast differentiation, and instead support a progressive remodelling imbalance in which an early low-turnover state is followed by increased osteoclast resorption.

The decrease in *Dlx5* and *Dlx6* at P35 is of particular interest in light of previous work linking Dlx genes to skeletal homeostasis and osteoblast–osteoclast coupling. Although the involvement of MeCP2 in the regulation of *Dlx5/Dlx6* has been debated, several studies have identified *Dlx5* as a potential MeCP2-responsive gene modulated by epigenetic mechanisms during differentiation [31–33]. Moreover, we previously showed that loss or reduction of *Dlx5* enhances osteoclast formation and activity without major changes in the Rankl/Opg ratio [30,34]. In this context, the early decrease in *Dlx5* and *Dlx6* observed here may reflect an upstream perturbation that contributes to the subsequent remodelling imbalance, even if the later osteoclast phenotype is not simply explained by changes in classical coupling markers.

The phenotype of the cortex of *Mecp2*^-/yBIRD^ long bones is also noteworthy. We observed an age-related reduction in marrow and external diameters associated with decreased cortical thickness. Because cortical bone is a major determinant of mechanical strength, these alterations are likely to contribute substantially to bone fragility. This progressive cortical impairment is consistent with the clinical observation that fractures and skeletal complications in RTT worsen with age. Although reduced periosteal apposition may contribute to this phenotype, the qPCR data do not indicate major alterations in late osteoblast differentiation markers at P55. It therefore remains possible that early regulatory defects, including the transient reduction in *Dlx5*/*Dlx6*, or altered osteoclast activity at the endosteal surface participate in cortical thinning.

An important point to consider is that the *Mecp2*^-/yBIRD^ mouse represents a severe loss-of-function model, whereas most patients with RTT carry specific *MECP2* mutations associated with variable residual protein function and heterogeneous clinical severity. Consistent with this, clinical studies indicate that skeletal manifestations are not uniform across RTT individuals, and that bone fragility is more pronounced in patients carrying specific mutations associated with greater disease severity, such as the nonsense R168X or R270X mutations. These observations suggest that the relative contribution of impaired bone formation and increased osteoclast resorption may vary depending on mutation type and disease progression. In this context, our findings likely reflect mechanisms associated with severe MECP2 deficiency and raise the possibility that osteoclast involvement may be particularly relevant at later stages or in more severe genotypes [13].

Overall, our findings refine the current model of skeletal involvement in RTT. Rather than reflecting only reduced bone formation, osteopenia in Mecp2 deficiency appears to result from a progressive remodelling imbalance, with an early low-turnover phase followed by increased osteoclast resorptive activity at later stages. Analysis of a public human osteoclast differentiation RNA-seq dataset (GSE246769; NCBI Gene Expression Omnibus) indicates that MECP2 is expressed throughout osteoclastogenesis, supporting the plausibility of cell-autonomous effects in this lineage. These results identify osteoclasts as important contributors to RTT-associated bone pathology and suggest that the skeletal phenotype evolves over time, a point that may be relevant for the design of future therapeutic strategies. In this context, a recent retrospective clinical study in females with RTT reported reduced lumbar BMD and provided preliminary evidence that zoledronate treatment may improve BMD in a subset of patients [35]. Given that zoledronate is a potent anti-resorptive bisphosphonate whose effects are mediated largely through inhibition of osteoclast differentiation/function and promotion of osteoclast apoptosis [36], this observation is consistent with the clinical relevance of increased osteoclast activity in RTT-associated bone fragility.

## ACKNOWLEDGEMENTS

We thank Corinne Collet (AH-HP department of molecular biology and biochemistry, Lariboisière hospital, Paris) for the measurements of DPD/Cr.

N.S. has been recipient of a doctoral fellowship from the French Ministry of Research. This research was partially supported by the EURORETT project and by the ANR project “DROS” and the EU Consortium IDEAL (HEALTH-F2-2011-259679) by Direction de la Recherche Clinique, Assistance publique-Hôpitaux de Paris.

